# Comparative Genomics Shows Differences in the Electron Transport and Carbon Metabolic Pathways of *Mycobacterium africanum* relative to *Mycobacterium tuberculosis* and suggests an adaptation to low oxygen tension

**DOI:** 10.1101/792762

**Authors:** Boatema Ofori-Anyinam, Abi Janet Riley, Tijan Jobarteh, Ensa Gitteh, Binta Sarr, Tutty Isatou Faal-Jawara, Leen Rigouts, Madikay Senghore, Aderemi Kehinde, Nneka Onyejepu, Martin Antonio, Bouke C. de Jong, Florian Gehre, Conor J. Meehan

**Affiliations:** Mycobacteriology Unit, Institute of Tropical Medicine, Antwerp, Belgium; Vaccines and Immunity Theme, Medical Research Council Unit, The Gambia, Banjul, The Gambia; Center for Global Health Security and Diplomacy, Ottawa, Canada; Department of Biomedical Sciences, Antwerp University, Antwerp, Belgium; Department of Medical Microbiology & Parasitology, University College Hospital, Ibadan, Nigeria; Department of Medical Microbiology & Parasitology, College of Medicine, University of Ibadan, Ibadan, Nigeria; Center for Tuberculosis Research, Nigeria Institute of Medical Research, Lagos, Nigeria; Division of Infectious and Tropical Diseases, London School of Hygiene & Tropical Medicine, London, United Kingdom; Medical School, University of Warwick, Coventry, United Kingdom; Bernhard-Nocht Institute for Tropical Medicine, Hamburg, Germany; School of Chemistry and Biosciences, University of Bradford, Bradford, United Kingdom

**Keywords:** Tuberculosis, *Mycobacterium africanum*, *Mycobacterium tuberculosis*, genome, Electron Transport, Carbon Metabolism

## Abstract

The geographically restricted *Mycobacterium africanum* lineages (MAF) are primarily found in West Africa, where they account for a significant proportion of tuberculosis. Despite this phenomenon, little is known about the co-evolution of these ancient lineages with West Africans. MAF and *M. tuberculosis* sensu stricto lineages (MTB) differ in their clinical, in vitro and in vivo characteristics for reasons not fully understood. Therefore, we compared genomes of 289 MAF and 205 MTB clinical isolates from the 6 main human-adapted *M. tuberculosis* complex lineages, for mutations in their Electron Transport Chain and Central Carbon Metabolic pathway in order to explain these metabolic differences. Furthermore, we determined, in silico, whether each mutation could affect the function of genes encoding enzymes in these pathways.

We found more mutations with the potential to affect enzymes in these pathways in MAF lineages compared to MTB lineages. We also found that similar mutations occurred in these pathways between MAF and some MTB lineages.

Generally, our findings show further differences between MAF and MTB lineages that may have contributed to the MAF clinical and growth phenotype and indicate potential adaptation of MAF lineages to a distinct ecological niche, which we suggest includes areas characterized by low oxygen tension.

## 1 Introduction

The *Mycobacterium tuberculosis* complex (MTBC) consists of a group of human–adapted ecotypes-*Mycobacterium tuberculosis* sensu stricto, *Mycobacterium africanum*, *Mycobacterium canetti* and animal-adapted ecotypes (1–4). There are seven known MTBC lineages (L) associated with particular geographic regions and adapted to specific human populations. These are the five lineages that make up *M. tuberculosis* sensu stricto (lineages 1-4 and lineage 7) and the two *M. africanum* lineages (lineages 5 and 6). Africa uniquely has a representation of all seven lineages.

MTBC strains from the seven lineages differ on average by about 1200 single nucleotide polymorphisms (5), with clear distinction between MAF and MTB lineages (6–8).

MTB Lineages 2 and 4 are more widespread geographically, more pathogenic and more transmissible, while MAF Lineages are exclusively found in West Africa and less transmissible (5, 9, 10). Clinically, MTB L4 is relatively more virulent than MAF L6 as evidenced by significantly faster progression, in contacts of infectious cases, to active disease (11). MAF lineages are associated with extrapulmonary disease and MAF L6 more commonly causes disease in immunocompromised persons and those with lower Body Mass Index, implying a more opportunistic pathogen (10, 12). Furthermore, MAF L5 and L6 grow markedly slower than MTB and prefer microaerobic growth conditions (13–15). Reasons for these differences are not completely known, although ours and other’s previous studies have attempted to explain some of the observations. A study documented non-synonymous SNPs or frameshift mutations in some genes associated with growth attenuation in MAF and higher mutation frequency in genes necessary for transport of sulphur, ions and lipids/fatty acids across the cell membrane (14). Another reported under-expression of, and MAF L6 specific mutations in, dormancy regulon genes, a network of genes crucial for the survival of MTB during hypoxia or anaerobiosis (15, 16).

Genes that encode proteins involved in nutrient metabolism and respiration, which are closely linked and together govern bacterial growth and survival, and those of unknown function, the conserved hypotheticals, are highest among the 4173 MTB H37Rv genes reported, emphasizing the importance of the nutrient metabolic and respiratory pathways (17).

As obligate aerobes, the mycobacteria respire and produce energy from varied nutritional or energy sources, such as carbon sources (18, 19). These enable the bacteria even to maintain metabolism without growth. The nutritional demands of the mycobacteria have been a topic of interest for over 100 years. Pioneering work throughout the 20th century elegantly showed that mycobacteria had unique nutritional requirements (20–29). Central to these findings was that different members of the MTBC were supported by different nutritional sources and consistent results of multiple phenotypic studies led to differentiating members of the MTBC based on their nutritional requirements (20, 30). For instance, it was observed that MTB showed eugonic growth on glycerol, while colonies of MAF and *M. bovis* were dysgonic, indicating an inability to properly utilize this carbon source. MAF and *M. bovis* were only able to show luxurious growth in the presence of sodium pyruvate (20, 30). Furthermore, MAF was found to grow small umbilicated colonies, which on paraffin embedded thin sections revealed extension deep into the media, rather than the surface (31).

Almost all the energy used by the bacteria is derived from the Central Carbon Metabolic Pathway. After nutrients are metabolized through this pathway, reducing equivalents are generated that eventually enter into the Electron Transport Chain for the generation of significant amounts of Adenosine Triphosphate (ATP) (18, 32).

Mycobacteria generate ATP via substrate level phosphorylation and oxidative phosphorylation, which produces more ATP through the activity of the F_1_-F_0_ ATP synthase in the Electron Transport Chain. Substrate level phosphorylation alone is insufficient to support growth of these bacteria (18).

Genome sequencing analyses show that the mycobacteria possess a branched respiratory pathway for electron transfer from electron donors to acceptors under different growth conditions. However, it appears that the transfer of electrons to oxygen, which seems to be the most preferred terminal electron acceptor, occurs with little plasticity, given that only two terminal oxidases have been found in mycobacteria, the bioenergetically more efficient *aa3*-type cytochrome c and the less efficient cytochrome *bd*-type menaquinol oxidases (33). During hypoxia or anoxia, mycobacterial growth is inhibited, even in the presence of alternate terminal electron acceptors within the branched respiratory chain such as fumarate and nitrate reductase. However, mycobacteria are still able to adapt and maintain metabolic functions (33).

Therefore, an intact Central Carbon Metabolic pathway and Electron Transport Chain are essential to mycobacterial growth and survival.

The importance of comparative genomics in unravelling the basis of distinct metabolic phenotypes in the MTBC has been shown. Using molecular genomic approaches, previous authors found that the inability of *M. bovis* to use glycerol and carbohydrates as sole carbon sources and its requirement for pyruvate in growth media was caused by a single nucleotide polymorphism in the *pykA* gene, encoding pyruvate kinase, resulting in a Glu220Asp amino acid substitution and causing the disruption of sugar catabolism (34). This mutation was also found in 3 MAF strains tested in the same analysis. Additionally, the authors showed that a frameshift at codon 191 of the *glpK* gene of the same *M. bovis* strain led to an incomplete coding sequence and the inability to use glycerol, although this *glpK* mutation was not present in all *M. bovis* strains.

Therefore, we aimed to investigate the genes involved in central carbon metabolism and respiration in the MTB and MAF lineages using a whole genome sequencing approach coupled with comparative genomics. We find important differences between the MAF and MTB lineages in their energy and nutrient metabolic pathways that likely contributed to the phenotypic differences observed between these lineages.

## 2 Materials and Methods

### 2.1 Ethical Statement

The study was conducted within the framework of an intervention trial of Enhanced Case Finding (ECF) in the Greater Banjul Area of The Gambia (Clinicaltrials.gov NCT01660646), piloted in 2012 and conducted between 2013 and 2017. This study was carried out in accordance with the recommendations of the Joint Gambia Government/MRC Ethics Committee and the Institute of Tropical Medicine, Antwerp Institutional Review Board. The protocol, including bacterial sub-studies, was approved by the Joint Gambia Government/MRC Ethics Committee and the Institute of Tropical Medicine, Antwerp Institutional Review Board. Nigerian isolates were collected from Southwest Nigeria within the West African Node of Excellence for TB, AIDS and Malaria (WANETAM) with the recommendations of the University of Ibadan and University College Hospital, Ibadan Joint Ethical Review Committees and the Nigerian Institute of Medical Research, Institutional Board (35). All subjects gave written informed consent in accordance with the Declaration of Helsinki and were anonymized.

### 2.2 Bacterial isolates

In The Gambia, MTB L4 followed by MAF L6 are the most isolated MTBC lineages. For almost a decade, the prevalence of all the MTBC lineages isolated in The Gambia has remained constant at 4.3% (L1), 2.5% (L2), 0.8% (L3), 57.2% (L4), 1.0% (L5), and 35.4% (L6) (36).

Within the framework of the ECF study conducted in the Greater Banjul Area, we sequenced 280 MAF L6 (32%), 3 MAF L5 (0.3%), 19 MTB L1 (2.2%), 36 MTB L2 (4%)(Beijing), 10 MTB L3 (1%) and 534 (60.5%) MTB L4 consisting of 85 MTB L4 Cameroon (9.6%), 15 MTB L4 Ghana (1.7%), 224 MTB L4 Haarlem (25.3%), and 211 MTB L4 LAM (23.9%). Given that only 3 MAF L5 were isolated from The Gambia within the period of analysis, we included 6 MAF L5 from Nigeria (Eastern West Africa), to improve the representativeness of this lineage.

Of this dataset, we analyzed the whole genome sequences of all 280 MAF L6 strains, the 3 MAF L5 isolated from The Gambia and the 6 MAF L5 from Nigeria, resulting in a total 289 MAF strains. For the MTB lineages, we analyzed a total 205 strains consisting of all 19 MTB L1, all 10 MTB L3 and 15 MTB L4 Ghana, 35 MTB L2 and a random number of L4 Haarlem (44), Cameroon (36) and LAM (46), while ensuring that all MTB L4 sublineages isolated were represented and isolates from each year of isolation, 2012 to 2014, were included.

### 2.3 DNA Extraction

Genomic DNA was extracted from loopfuls of pure MTBC colonies grown on Lowenstein-Jensen media (37) using the Maxwell 16 DNA Purification Kit (Promega). DNA from Nigerian strains was extracted using the Cetyl trimethylammonium bromide (CTAB) method (38).

### 2.4 Whole-genome sequencing

Sequencing of MTBC isolates was performed at MicrobesNG, Birmingham; GenoScreen, France; FISABIO, Valencia or the Beijing Genome Institute (BGI), Beijing. Sequencing reads were generated on a HiSeq or Miseq platform (IIIumina). Quality control was performed for each provider to ensure adequate sequencing depth (>30X) and genome coverage (>95% of the H37Rv reference strain). Raw Illumina reads have been deposited in the ENA with accession <to be undertaken on acceptance>.

### 2.5 Bioinformatics Analysis

#### 2.5.1 Mapping and Variant calling

We used Snippy version 3.1 for the analysis of genomes. Briefly, paired-end raw reads of each sample were mapped to the *M. tuberculosis* H37Rv reference genome (GenBank accession number: NC_000962.3) using BWA-MEM 0.7.12 (39). Mapped reads were converted to the SAM/BAM format and sorted using Samtools 1.3 1 (40). Variant calling was done using Freebayes 0.9.20 (41). Variants were called only if ≥10 reads covered variant positions and ≥90% of those reads differed from the reference. Genes were annotated with SnpEff 4.1 (42). Samples were assigned to MTBC lineages based on the classification of Coll and colleagues (43) using the VCF output of snippy and the PhyResSE SNP list (44).

#### 2.5.2 Phylogenetic analysis

Using the Snippy output folders of all isolates and custom python scripts, a SNP alignment and a count of all excluded invariant sites were created. A maximum-likelihood (ML) phylogeny was inferred using RAxML version 8.2.9 (45) executing a thousand rapid bootstrap inferences under the general time-reversible (GTR) model, with ascertainment bias correction using the Stamatakis reconstituted DNA approach (46, 47). The resulting tree was exported to interactive Tree of Life (iTOL) for visualization (48).

#### 2.5.3 Protein Predictions

The effect of amino acid substitutions were predicted using PROVEAN software tools at default settings (PROVEAN protein) (49–51). The PROVEAN NCBI non redundant 2012 database is a newer and larger sequence database than SIFT databases (52) but comparable to SIFT in prediction accuracy. A PROVEAN score of ≤ −2.5 implies that the amino acid substitution could impact negatively on protein function.

## 3 Results and Discussion

Given that generalists and specialists differ in the vastness of their ecological niches (53), and thus carbon sources, we compared the specialist MAF lineages (289 strains) to the generalist MTB lineages (205 strains) for genomic differences in their Electron Transport and Carbon metabolic pathways. These pathways are intrinsically linked and are central to deriving energy (ATP) from carbon sources (Figure 1).

**Figure 1:**
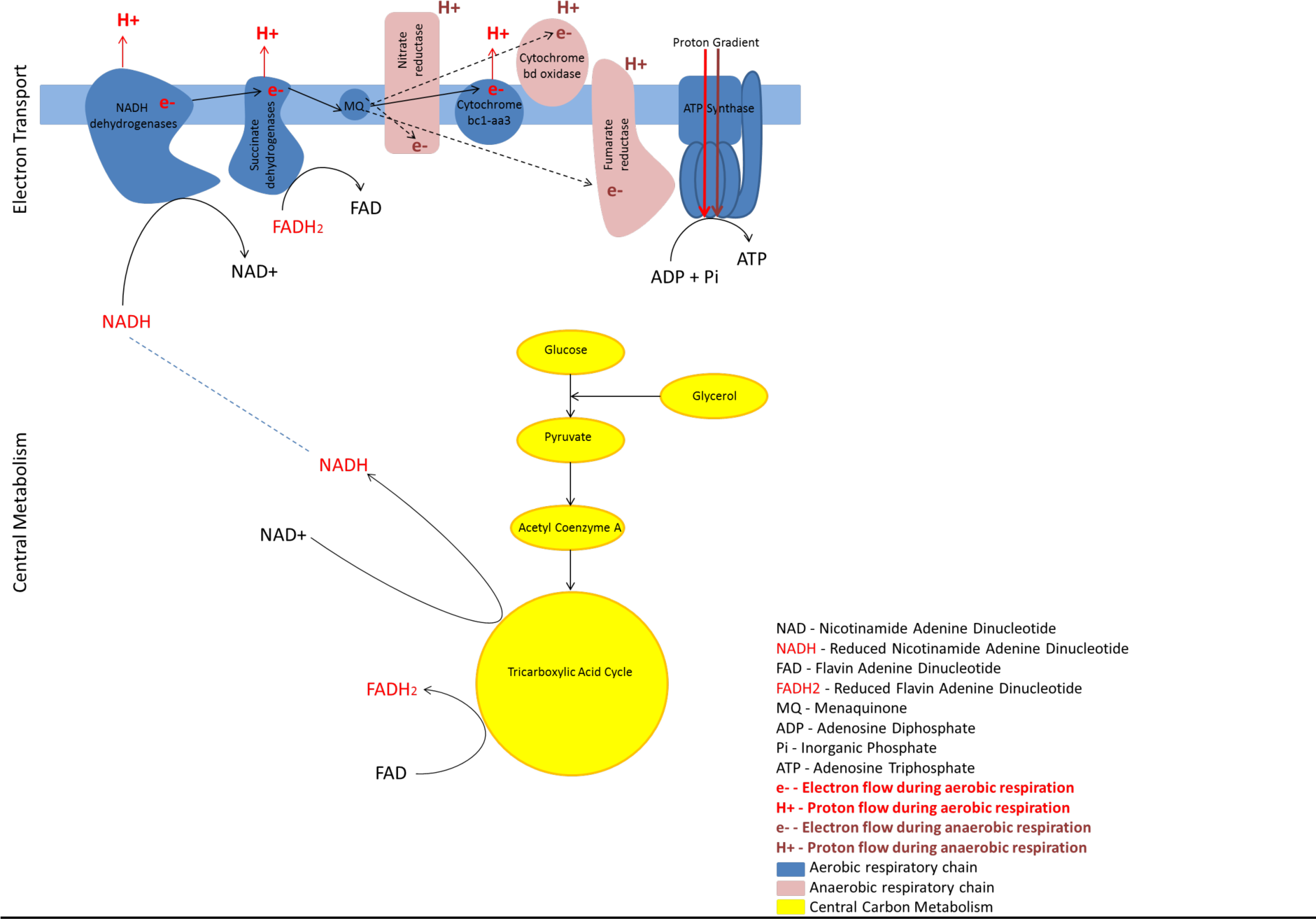
Link between central carbon metabolism and the Electron Transport Chain. Reducing equivalents like NADH and FADH2 enter into the Electron Transport Chain that drives the production of high amounts of ATP for cellular processes

Understanding differences in metabolism and respiration between the MTBC lineages is also pertinent for tackling TB as mycobacterial central metabolism and respiration have re-emerged as potential targets for TB chemotherapy. The new TB drug, Bedaquiline (TMC207) and the drug candidate, Telacebec (Q203), both target the respiratory chain (54–56) and even the repurposed drug Clofazimine, and another new drug, Delamanid, reportedly interfere with redox cycling and cellular respiration by generating Reactive Intermediates (57–61). Thus respiratory inhibitors may offer the next generation of core drugs against mycobacterial diseases (62) and there is a pressing need to understand the differences in these processes between the different lineages of the MTBC.

It is now increasingly apparent that genetic and metabolic differences between the MTBC human-adapted lineages have the potential to affect transmission, diagnostics and treatment (5, 63–68). Moreover, niche adaptation is influenced by the metabolic requirements of an organism, a consequence of evolution. However, the extent of such genetic changes and their potential metabolic effects has not been well explored. In this study, we found a large number of mutations with the potential to negatively affect gene function (hereafter referred to as harmful mutations), particularly in MAF lineages (Figure 2 and Figure 4). However, we also observed that similarities in these pathways occurred between some MTB and MAF lineages potentially due to convergent evolution.

**Figure 2:**
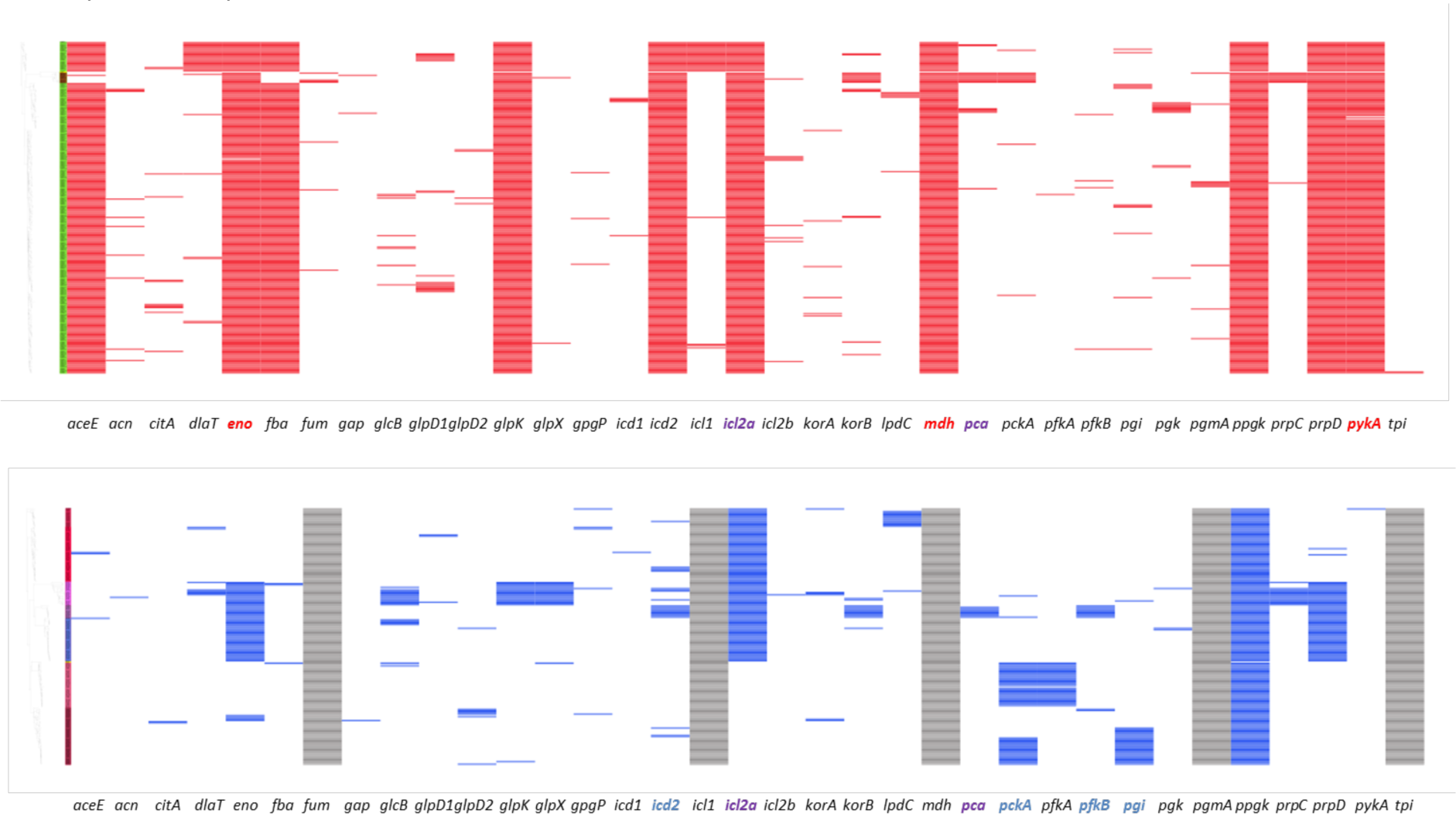
Mutations detected in the Central Carbon Metabolic Pathway of a) MAF L6 and MAF L5 and b) MTB L4 Ghana, L4 Haarlem, L1, L3, L2, L4 Cameroon and L4 LAM. Red and blue coloured bars represent nonsynonymous mutations detected in 50-100% of strains per lineage in the associated genes shown below each set of bars indicating mutations. Genes emboldened red have mutations detected predicted to be harmful in MAF lineages, genes emboldened blue have mutations detected predicted to be harmful in MTB lineages and genes emboldened purple have mutations detected predicted to be harmful in the same gene in MTB and MAF lineages. Grey bars indicate that no nonsynonymous and/or predicted harmful mutations were detected in the corresponding gene below each set of grey bars. First column colour indicates the lineage: a) MAF L6 (green) and MAF L5 (brown), b) MTB L4 Ghana (brown), L4 Haarlem (red), L1 (lilac), L3 (purple), L2 (blue), L4 Cameroon (pink) and L4 LAM (dark red).

### 3.1 Mutations in genes encoding Central Carbon Metabolic pathway enzymes

We examined mutations in all genes in the glycolytic pathway, the Tricarboxylic Acid cycle and the Methylcitrate Cycle. Genes and enzymes in these pathways contribute directly to the growth and virulence of the MTBC, by generating products that feed directly into the Electron Transport Chain (69). Therefore, mutations, with the potential to affect proper function of these genes and their gene products, would have obvious consequences downstream, for energy generation, survival and ultimately, the virulence of members of the MTBC.

#### 3.1.1 The Glycolytic Pathway

In MAF lineages, all genes leading to the breakdown of sugars to 2-phosphoglycerate at the seventh step (Figure 2 and Figure 3), were generally conserved. However, at the critical stage of pyruvate metabolism, mutations with the potential to affect the normal function of Enolase (*eno*), Pyruvate kinase (*pykA*) and Pyruvate carboxylase (*pca*) were detected. Enolase is responsible for the penultimate step of glycolysis, where 2-phosphoglycerate is converted to Phosphoenolpyruvate. Pyruvate kinase modulates the irreversible reaction converting Phosphoenolpyruvate to Pyruvate while Pyruvate carboxylase drives the conversion of Pyruvate to Oxaloacetate. Of these three genes, it had previously been shown, in *M. bovis* and 3 MAF strains, that the mutation in *pykA* rendered the critical enzyme pyruvate kinase inactive, resulting in a metabolic block at the level of pyruvate biosynthesis (34). We also confirm this mutation in the MAF strains we analyzed in our study (Figure 2). However, to the best of our knowledge this is the first report on lineage-specific mutations with potentially harmful effects in the essential gene *eno*, in MAF L6. *eno* is found on the surface of many pathogenic bacteria and beyond its primary role in glycolysis, aids in tissue remodeling and invasion of host cells (70). This gene was recently reported as a novel target for the 2-aminothiazoles of the aminothiazole (AT) series, that exerted its effect by inhibiting *eno* in MTB H37Rv (L4 strain). Therefore, the potentially harmful mutations we detected in *eno* may affect energy metabolism and potentially contributes to the metabolic block at the level of pyruvate biosynthesis, further reducing the fitness of L6.

**Figure 3:**
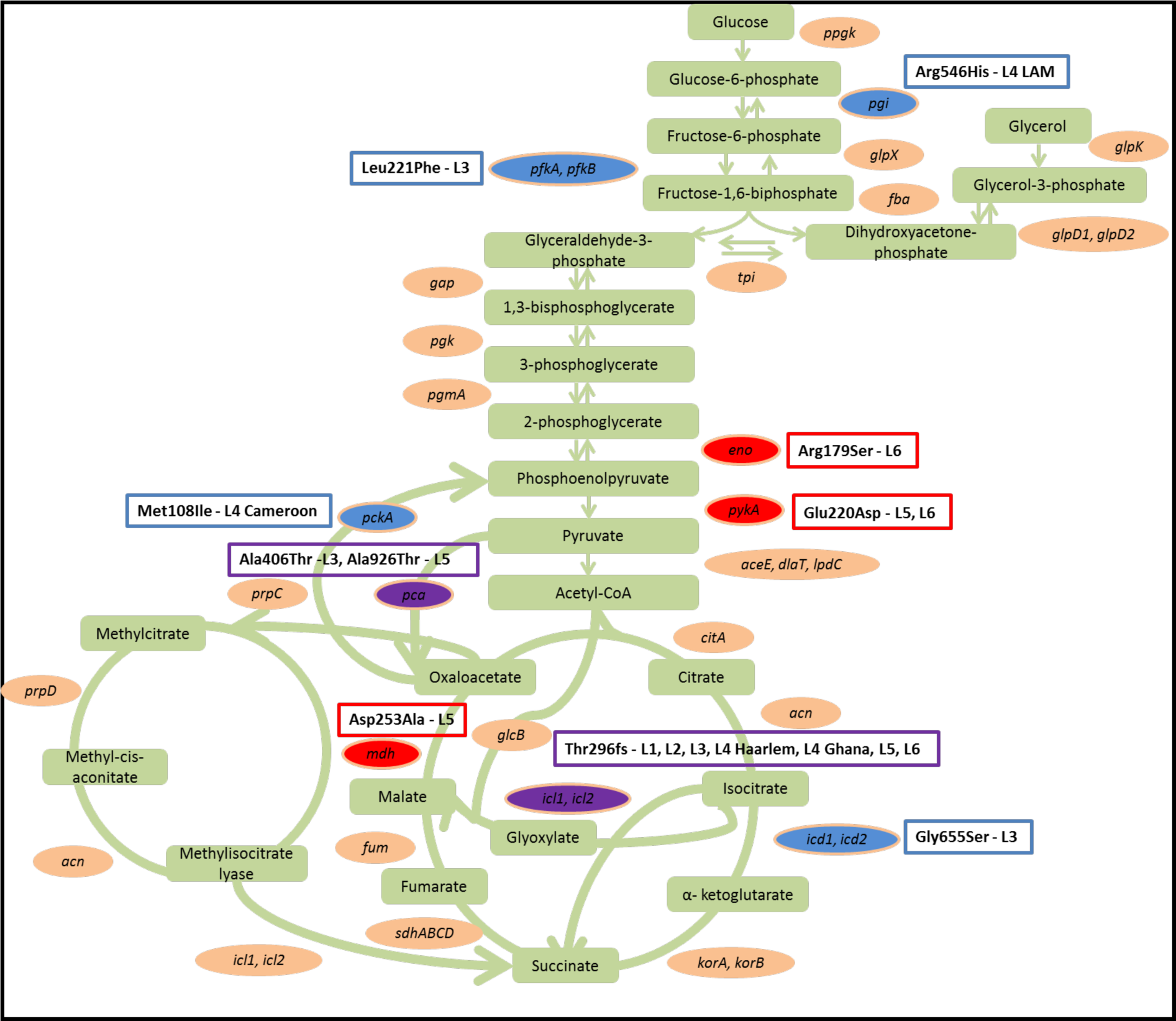
Mutations in the Central Carbon Metabolic Pathway of MAF and MTB. Genes with a **red** colour code have harmful mutations in MAF lineages, indicated next to the gene. Genes with a **blue** colour code have harmful mutations in MTB lineages, while genes in **purple** have harmful mutations in the same gene in MAF and MTB lineages.

Unlike the MAF lineages, MTB lineages, notably, did not have any potentially harmful mutations in *eno* and *pykA* (Figure 2 and Figure 3), implying a general ability to complete glycolysis. However, mutations were detected in Glucose-6-phosphate isomerase (*pgi*) in L4 LAM and in Phosphofructokinase B (*pfkB*) in MTB L3, at the second and third steps of glycolysis respectively, where Glucose-6-phosphate is converted to Fructose-6-phosphate and subsequently to Fructose-1-6-biphosphate (Figure 3). *pgi* is essential for the in vitro growth of *M. tuberculosis*, and *M. smegmatis pgi* mutants are glucose auxotrophs. Moreover, an additional role for *pgi* in cell wall biosynthesis was reported (37, 71–73). Therefore, *pgi* is important and the effect of the predicted harmful mutations we detected in *pgi* in L4 LAM should be investigated. However, the defect in *pfkB* in MTB L3 is unlikely to affect glycolysis given that Phosphofructokinase activity has so far only been associated with *pfkA*, where a *pfkA* deletion mutant was neither able to grow on glucose in vitro nor to have any detectable Phosphofructokinase activity; the mutant could not be rescued by expressing *pfkB* (32, 74).

Interestingly, additional genes, Pyruvate carboxylase (*pca*) in MAF L5 and MTB L3 and Phosphoenolpyruvate carboxylase (*pckA*) in MTB L4 Cameroon, were mutated at the level of pyruvate metabolism (Figure 3). *pca* and *pckA* secrete enzymes that contribute significantly to the control of metabolic flux to glycolysis, gluconeogenesis and anaplerosis. These genes carry out functions related to cholesterol detoxification and lipogenesis during intracellular survival and *pckA* was shown to be essential for virulence in *M. bovis* (75, 76).

#### 3.1.2 The Tricarboxylic Acid Cycle

At the final stage of glycolysis, pyruvate is generated and converted to Acetyl Coenzyme A (Figure 1 and Figure 3), that is fed into the critical Tricarboxylic Acid Cycle (TCA), for the release of reducing equivalents into the Electron Transport Chain. Therefore, the ability to complete this cycle, whether by progression through all stages of the cycle or via the crucial glyoxylate by-pass, is essential for the generation of ATP.

Overall, genes encoding enzymes of the TCA cycle were largely conserved in MAF and MTB lineages, however, in MTB L1-L3, L4 Haarlem, L4 Ghana and MAF lineages, we found potentially harmful mutations in Isocitrate lyase (*icl2a*) and Isocitrate dehydrogenase (*icd2*).

Emphasizing the importance of the glyoxylate shunt, *M. tuberculosis* strains lacking both ICLs are unable to grow on fatty acids in vitro, establish and maintain a chronic infection in mice and were said to be the most severely attenuated strains (32, 77). In our analysis, we found the frameshift mutation previously found in H37Rv, an L4 strain (78), which we, for the first time to the best of our knowledge, also report in MTB L4 Haarlem and Ghana, MTB L1, L2, L3, MAF L5 and L6, yet, interestingly, not in MTB L4 Cameroon and LAM (Figure 2, Figure 3 and Table 1).

**Table 1:** Potentially deleterious mutations in genes encoding enzymes in the Central Carbon Metabolic Pathway. List of all mutations found to be potentially harmful by PROVEAN (marked by a *) are listed. The full list of mutations is given in supplementary table S5.

| Gene | MTB amino acid change | MTB Lineage | MAF amino acid change | MAF Lineage |
| --- | --- | --- | --- | --- |
| *pgi* * | Arg546His | clade of L4<br>LAM |  |  |
| *pfkB* * | Leu221Phe | L3 |  |  |
| *eno* * | Arg8Gly<br>Lys429Gln | L1, L2, L3 | Arg8Gly<br>Arg179Ser | L5, L6 |
| *pykA* * |  |  | Glu220Asp | L5, L6 |
| *pca* * | Ala406Thr | L3 | Ala926Thr | L5 |
| *icd2* * | Gly655Ser<br>Leu254Val | L3 | Val447Met<br>Lys117Asn | L6 |
| *mdh* * |  |  | Leu326Ile<br>Ala112Thr<br>Asp253Ala | L5, L6 |
| *icl2a* * | Gly179Asp<br>Thr296fs | L1,L2,L3,L4<br>Haarlem, L4<br>Ghana | Ala146Thr<br>His151Arg<br>Gly179Asp<br>Thr296fs | L5, L6 |
| *pckA* * | Met108Ile<br>Ala536Gly | L4<br>Cameroon,<br>clade of L4<br>LAM | Lys422Thr | L5 |

As stated earlier, in section 3.1.1, Pyruvate carboxylase (*pca*), that converts pyruvate to oxaloacetate towards the final stage of glycolysis, was mutated in L5, however, at the last step of the TCA cycle, malate dehydrogenase (*mdh*) that converts malate to oxaloacetate, was also mutated (Figure 2, Figure 3 and Table 1), implying that both routes for the production of oxaloacetate in MAF L5 have mutated genes. Given that oxaloacetate feeds not only into the TCA cycle but also into the methylcitrate cycle, impaired function of *mdh* may affect energy metabolism in MAF L5.

Overall, with major blocks in Central Carbon Metabolism in MAF lineages, the number of reducing equivalents produced via the central carbon metabolic pathway for electron transport may be lower. MTB lineages largely had a conserved glycolytic pathway and TCA cycle, however, different MTB lineages have previously been shown to have different growth rates and patterns (79–81). Notably, the growth rate of MTB L3 is reportedly lower compared to other MTB lineages (79, 80). Therefore, in MTB lineages/sublineages where we found more potentially harmful mutations, particularly in MTB L3, further investigations on the effect of these mutations on energy metabolism need to be carried out as slower growth of these lineages may be directly linked to impaired energy metabolism.

### 3.2 Mutations in genes encoding Electron Transport Chain enzymes

As reducing equivalents, like NADH, from the Central Carbon Metabolic Pathway deliver electrons into the Electron Transport Chain, NADH dehydrogenases serve as the gateway of the Electron Transport Chain in the MTBC and electrons are transferred from NADH oxidation to quinone reduction and ultimately to ATP synthase for ATP production in greater quantities (Figure 1). Therefore, defects in the Electron Transport Chain are bound to affect the net yield of ATP generated.

Relative to MTB lineages, MAF lineages had multiple mutations predicted to affect the normal function of genes encoding key enzymes of the Electron Transport Chain (Figure 4 and Figure 5). Potentially harmful mutations were detected in *ndhA*, *pruB*, *qcrC*, *ctaB*, *frdA*, *atpHG*, *sdhA* and *ald* (Figure 4, Figure 5, Table 2 and Supplementary File S3). Interestingly, for some MTB lineages, mutations predicted to affect gene function were also found in *ndhA*, *Rv0249c*, *frdB*, *atpD*, and *nuoDHF* (Figure 4, Figure 5, Table 2 and Supplementary File S4).

**Table 2:**
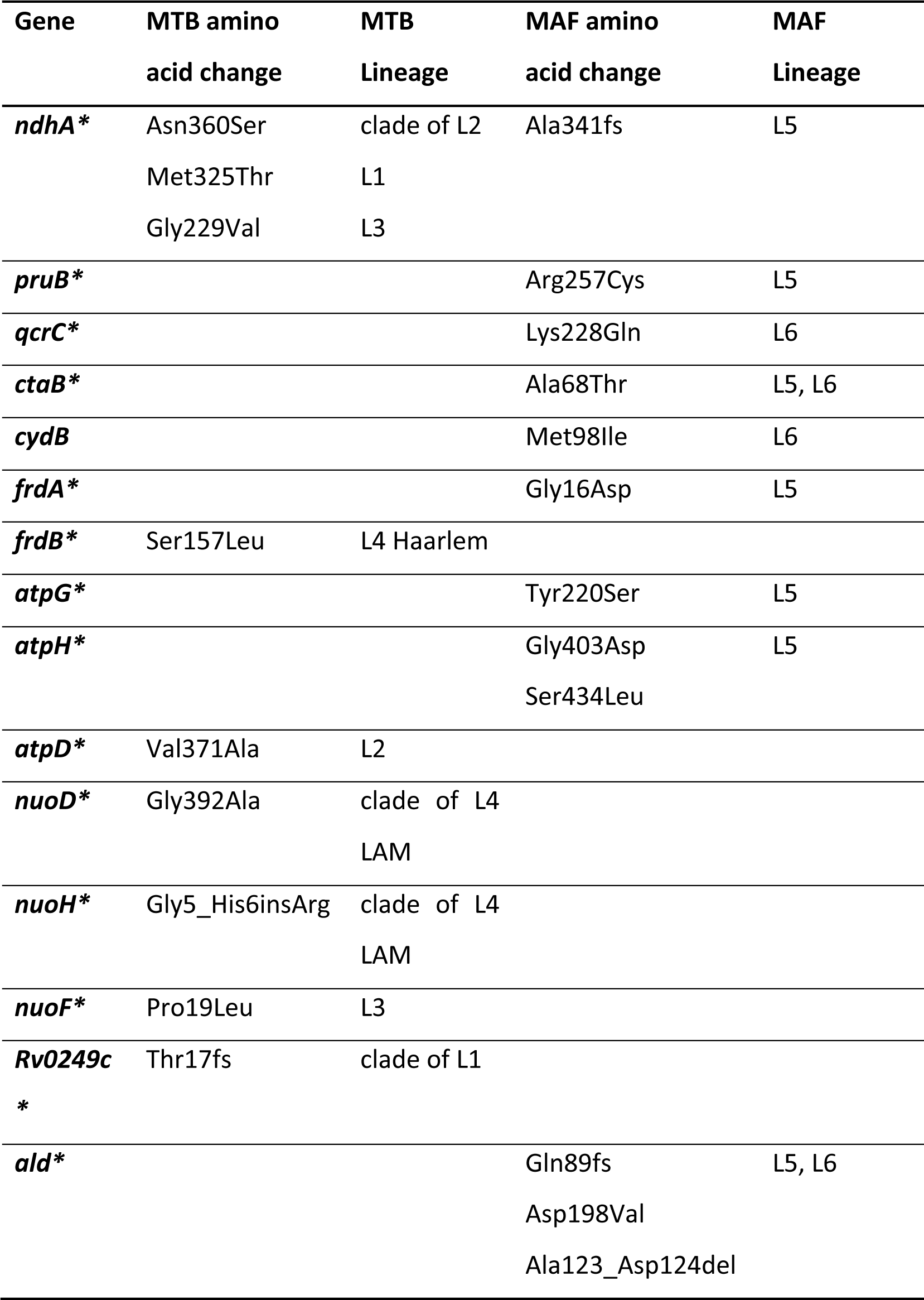
Potentially deleterious mutations in genes and gene subunits encoding Electron Transport Chain enzymes. List of all mutations found to be potentially harmful by PROVEAN (marked by a *). The full list of mutations is given in supplementary table S5.

**Figure 4:**
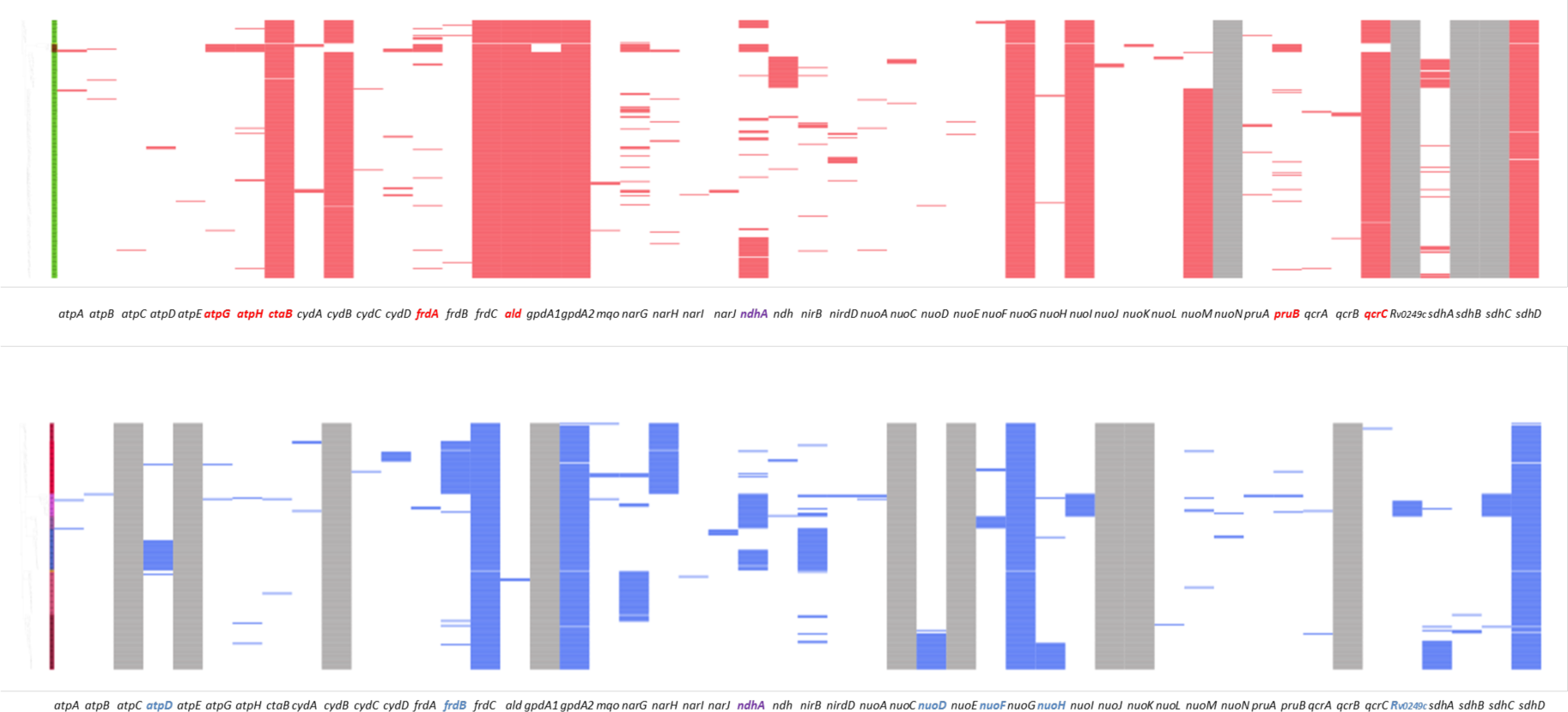
Mutations detected in the Electron Transport Chain of a) MAF L6 and MAF L5 and b) MTB L4 Ghana, L4 Haarlem, L1, L3, L2, L4 Cameroon and L4 LAM. Red and blue coloured bars represent nonsynonymous mutations detected in 50-100% of strains per lineage in associated genes shown below each set of bars indicating mutations. Genes emboldened red have mutations detected predicted to be harmful in MAF lineages, genes emboldened blue have mutations detected predicted to be harmful in MTB lineages and genes emboldened purple have mutations detected predicted to be harmful in the same gene in MTB and MAF lineages. Grey bars indicate that no nonsynonymous and/or predicted harmful mutations were detected in the corresponding gene below each set of grey bars. First column colour indicates the lineage: a) MAF L6 (green) and MAF L5 (brown), b) MTB L4 Ghana (brown), L4 Haarlem (red), L1 (lilac), L3 (purple), L2 (blue), L4 Cameroon (pink) and L4 LAM (dark red).

**Figure 5:**
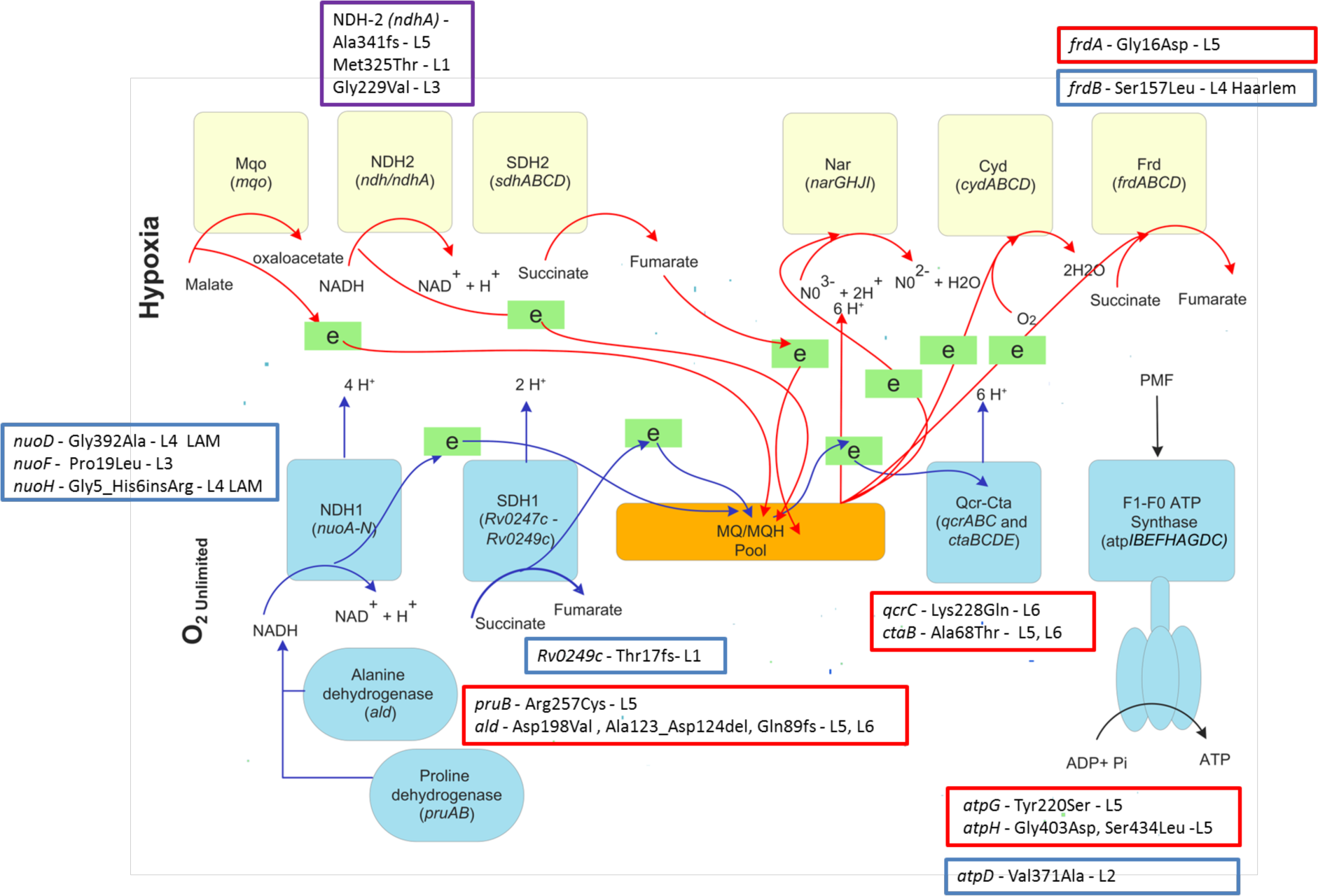
The core and alternate respiratory chain of mycobacteria during in vitro exponential growth when oxygen is abundant and when oxygen is limited. Mutations in MAF lineages are indicated in **red** boxes next to each affected complex. Mutations in MTB are indicated in **blue** boxes next to the affected gene while mutations found in the same gene in both MAF and MTB are shown in **purple** boxes. The core chain consists of type I NADH:menaquinone oxidoreductase (NuoA-N), succinate: menaquinone oxidoreductase 1 (SDH1), cytochrome aa3-bc supercomplex (Qcr-Cta) and F1F0-ATPase while the route used during oxygen limitation is composed of Type II NADH:menaquinone oxidoreductase (Ndh); succinate:menaquinone oxidoreductase (SDH2); nitrate reductase (Nar); Fumarate Reductase (Frd) and cytochrome bd oxidase (Cyd), a high affinity terminal oxidase allowing hypoxic survival.

#### 3.2.1 NADH dehydrogenases

To receive NADH from central metabolism into the Electron Transport Chain, *M. tuberculosis* possesses two NADH dehydrogenases, NDH-1 and −2. NDH-1 has 14 subunits (*nuo*A-N) while two copies of NDH-2 exist in *M. tuberculosis*: *ndh* and *ndhA*. Between the two NADH dehydrogenases, an essential role of *ndh* for the growth of *M. tuberculosis* was reported previously and a function of *ndh* in recycling NADH and maintaining an energized membrane was documented (33, 82). Even though *ndhA* was previously reported to be dispensable for growth, it was recently shown that both *ndh* and *ndhA* differentially control oxygen consumption (82). In fact, *ndh* was also shown to be dispensable for growth of *M. tuberculosis* but deletion of both *ndh* and *ndhA* prevented growth altogether in standard media and resulted in attenuated growth in mice (82, 83). We found a potentially harmful mutation in *ndhA* in all MAF L5. Given the adaptation of MAF lineages towards a microaerophilic lifestyle, *ndhA* may be redundant in MAF L5 and defects in this gene may have coincided with the adaptation to a microaerophilic lifestyle (84).

No potentially harmful mutations were detected in the essential *ndh* in any analyzed genome. This implies that the function of *ndh* is likely conserved across the MTBC lineages.

#### 3.2.2 Succinate dehydrogenases

Succinate dehydrogenase 1 (*Rv0247c-Rv0249c*, SDH-1) functions during aerobic respiration when oxygen and nutrients are abundant, while SDH-2 (*sdhABCD*) functions during hypoxia/anaerobiosis, when nutrients are limited (33, 85, 86) (Figure 5). Succinate dehydrogenases are a direct link between the TCA cycle and electron transport and typically reduce succinate to fumarate (87).

The only potentially harmful mutations we detected in the Succinate dehydrogenases were in *sdhA* of SDH2 in a clade of MAF L6 and in SDH1, *Rv0249c*, in a clade of MTB L1 (Figure 4). Given the essential role of SDH1 for growth and survival, where the deletion of SDH1 was shown to impair the rate of respiration through the Electron Transport Chain and to reduce cell viability (85), future studies should determine if electron transport and aerobic respiration are affected in certain MTB L1 strains, as recent studies report slower growth and a lower odds to grow in culture for MTB L1 and the MAF lineages (67, 79).

#### 3.2.3 Cytochrome bc1-aa3 complexes and Cytochrome bd oxidase

Electrons from Succinate dehydrogenases, move into the quinone pool, ready for transfer to the third and fourth complexes Cytochrome bc1-aa3 (*qcrABC* and *ctaBCDE*), during aerobic respiration, when oxygen is abundant, and to the less efficient cytochrome bd oxidase (*cydABCD*) when oxygen is limited (Figure 5).

The bc1-aa3 complexes are the major respiratory route in mycobacteria under standard aerobic conditions and are essential for growth where they play a key role in oxidative phosphorylation and electron transport that yields more ATP (33, 88), yet in our analysis, we detected potentially harmful mutations in *qcrC* and *ctaB* of the bc1-aa3 complexes in MAF lineages and intact gene complexes in MTB lineages (Figure 4 and Figure 5). To the best of our knowledge, this is the first work demonstrating that the critical bc1-aa3 complex is mutated in MAF lineages. This is perhaps crucial to understanding the difference in energy metabolism, particularly oxidative phosphorylation, between the MTB and MAF lineages, especially because MAF lineages preferentially grow microaerobically (15, 84, 89–91) and significantly under-express the dormancy regulon required for adaptation to oxygen limitation (15). Notably, the Imidazopyridine amide in Phase 2 clinical trials, Telacebec (Q203), inhibits *qcrB* of Cytochrome bc1, further emphasizing the importance of this complex for the survival of MTB. Therefore, the predicted harmful mutations we found in *qcrC*, the heme of the cytochrome bc1-complex, in MAF L6 (Figure 4 and Figure 5) likely impairs aerobic respiration and growth in this lineage severely, given that *qcrC*, like *qcrB*, is essential for survival (86). Similarly, *M. smegmatis* strains that had mutations in the bc1-aa3 complex were significantly growth impaired, confirming the essentiality of the bc1-aa3 respiratory pathway for mycobacterial growth. Recently, it was also confirmed that *M. tuberculosis* requires the bc1-aa3 complex to attain optimal growth rates and high titres in mice (83).

Taken together, our analysis provides further support for the view that MAF is adapted to a distinct niche, less dependent on aerobic respiration and more adapted to a microaerobic lifestyle (15). Reasons for this potential niche adaptation and the benefit to the pathogen in its interaction with its host should be investigated further.

Ultimately, with an impaired bc1-aa3 complex, ATP yield will be reduced overall. This is likely the case in MAF lineages. Therefore, we postulate that ATP yield through oxidative phosphorylation in the MAF lineages is lower compared to the MTB lineages.

#### 3.2.4 ATP Synthase

The F_1_-F_0_ ATP synthase itself, the target for Bedaquiline (54, 92), is rather conserved in the different lineages. However, in MAF L5 and MTB L2 we detected potentially harmful mutations in *atp* genes (Table 2). The F_1_-F_0_ ATP synthase operon is encoded by *atpIBEFHAGDC* and is required for survival as all genes in the operon are essential (71, 93). A defective ATP synthase may coincide with the microaerophilic lifestyle of MAF. However, for MTB L2, it is not clear what the advantage is for acquiring mutations in genes encoding ATP synthase. Interestingly though, compared to MTB L4, MTB L2 was found to have a lower growth rate (79).

#### 3.2.5 Fumarate reductase

The branched respiratory chain of *the* MTBC permits anaerobic survival. During hypoxic or anaerobic conditions when oxygen is limited, Fumarate reductase (FRD, *frdABCD*) and Nitrate reductase (NAR, *narGHJI*) can serve as terminal electron acceptors to maintain the membrane potential. Therefore, these enzymes are critical. Moreover, further studies of *sdh* suggested that FRD could partially compensate for a lack of SDH activity (94). Defects in any of the subunits of *frd* could limit the overall function of the FRD complex. Notably, the attenuated strain H37Ra grown under low-oxygen conditions showed a lag in gene expression of *frdA* and *frdB* (95). The mutations we found in *frdA* and *frdB* in MAF L5 may be related to the adaptation of this lineage to a hypoxic or microaerophilic lifestyle. Therefore, the effect of mutations on gene function in MAF L5 and MTB L4 sublineage, Haarlem, should be determined experimentally.

#### 3.2.6 Alanine dehydrogenase and Proline dehydrogenase

Other key dehydrogenases that contribute to redox balance of NADH for initiation of electron transport were also mutated in the MAF lineages (Figure 4 and Figure 5), indicating potential redox imbalance in MAF lineages with the likelihood to impede electron transport. From our analysis, all MAF lineages had potentially harmful mutations in Alanine dehydrogenase (*ald*) and all MAF L5 had potentially harmful mutations in Proline dehydrogenase (*pruB*). Proline dehydrogenase is associated with the adaptation to hypoxia, slow growth rate and is essential for growth (71, 93, 96, 97). Alanine dehydrogenase has been shown to play a role in redox balance during low oxygen conditions and the downshift of *M. tuberculosis* to the state of nonreplicating persistence. *ald* mutants had altered NADH/NAD ratios and significant delays in growth resumption after reaeration. Additionally, induction of *ald* rescued the bc1-aa3 complex mutant while its disruption made the growth defect of the mutant worse (97, 98).

Given that *ald* and the bc1-aa3 complex were mutated in all MAF in our analysis, MAF lineages are most probably natural mutants of *ald* and the bc1-aa3 complex. It is possible that these mutations in the MAF lineages contribute significantly to their slower growth compared to MTB due to impaired energy production. Moreover, these polymorphisms further support the adaptation of MAF to a hypoxic niche.

### 3.3 Additional Mutations and similarities between MTBC lineages in the Central Carbon Metabolic Pathway and Electron Transport Chain

We detected several other mutations in genes of the Central Carbon Metabolic pathway and Electron Transport Chain that could potentially impact on enzyme function (Supplementary File S5). However, their PROVEAN scores were above the cut-off for harmful mutations and thus any impact may be slight or only due to neutral evolution. *cydB* and *narG*, highly important genes in the Electron Transport Chain also had several mutations, but none predicted to be potentially harmful (Supplementary File S5).

For multiple genes in the Electron Transport Chain and the Central Carbon Metabolic pathway, we either found the same mutation occurring in the same gene or different mutations occurring in the same gene in the different MTBC lineages. These mutations may confer a selective advantage and/or contribute to adaptation (Supplementary File S5).

Of these, only those mutations detected in *pca* in MTB L3 and MAF L5, those detected in *icl2a* and those found in *ndhA* are potentially harmful.

Overall, more similarities occurred between MTB L1, MTB L3 and the MAF lineages. This is interesting as the MAF lineages, MTB L1 and L3 lineages all reportedly grow relatively slower than the MTB L2 and L4 lineages (14, 67, 79, 80, 99). We postulate that the slower growth rate of these lineages and the similarities we observe in their metabolic pathways, may be linked to their similar migration and dispersal patterns in Africa and Eurasia (100), where L2 and L4 have become widely dispersed, while L5, L6, and L7, had more geographically restricted expansion, adapting to more specific hosts. This may have influenced niche adaptation, where L2 and L4, in line with increased dispersion and range expansion, also increased their replicative/growth capacity and ability to transmit, while the more host restricted lineages maintained a lower replicative/growth potential in line with expansion in situ.

### 3.4 Limitations

Limitations of our analysis include the small sample size of MAF L5 analyzed. In the Gambia and other countries in Western West Africa, the prevalence of L5 is significantly lower than L6 (36). In our isolation of the 289 MAF strains included in this study, only 3 from The Gambia were MAF L5. However, to ensure that the mutations we detected in L5 from The Gambia were more representative of the MAF L5 lineage, we included L5 from Nigeria (Eastern West Africa), where the prevalence of L5 is high and L6 is significantly less likely to be isolated (101). Another limitation is that we did not experimentally confirm any of our in silico phenotype predictions.

## 4 Conclusion

In this comparative analysis, we describe genomic differences between the MTB and MAF lineages in genes encoding enzymes of the Electron Transport Chain and the Central Carbon Metabolic pathway, which may explain the differences in the clinical- and in vitro phenotype described for the MAF and MTB lineages. In vitro, MAF lineages grow significantly slower than MTB lineages and MAF L6 is, clinically, less virulent than MTB L4, as evidenced by significantly lower progression of MAF L6 infected individuals to active TB disease (11). Again, in vitro, MAF lineages show microaerobic growth and clinically, are associated with extrapulmonary disease, implying a preference for regions with low oxygen (10, 15). Furthermore, MAF L6 more commonly causes disease in immunocompromised persons, implying a more opportunistic pathogen (12, 13). Generally, it appears from our analysis that compared to MTB lineages, MAF lineages had the most mutated Central Carbon Metabolic and Electron Transport Pathways, with mutations occurring in critical components of each pathway. The combined effect of a defective Carbon Metabolic Pathway and Electron Transport Chain in MAF lineages, likely contributes to the reduced fitness of the MAF lineages. We speculate that our findings may contribute to 1 - the slower growth of MAF lineages, 2 - relative attenuation of the MAF L6 lineage compared to MTB lineages and 3 - host specificity to West Africans.

It is intriguing that in the different MTBC lineages multiple harmful mutations occurred in the same gene. These similarities largely found in MTB lineages 1, 2, 3 and the MAF lineages, indicates there may be a selective advantage for this. Interestingly, these lineages, compared to MTB L4 strains, have been associated with slower growth and cytokine induction patterns suggestive of immune evasion (8, 14, 67, 79–81, 99, 102, 103).

If the potentially harmful mutations we report on in this analysis affect energy metabolism in the different MTBC lineages, fitness will differ and ultimately the infection and transmission potential of these lineages. Therefore, our findings are not only relevant for TB product development but also for transmission studies and interventions. The literature already provides credence to and evidence for our hypothesis (9, 81, 104, 105).

### 4.1 Future Proposed Investigations

#### 4.1.1 Functional Studies

Significant genomic differences between MTB and MAF lineages presented in this analysis warrant further studies in order to properly characterize the regulatory network of MAF. Recently, the regulatory network of H37Rv (MTB L4) was characterized yet what pertains to MAF lineages is not clearly defined, even though it has been shown that master regulators like *PhoPR* and *Rv0081* as well as the DosR regulon are underexpressed and/or mutated in MAF (15, 106–108). It is possible that the PhoPR system controls other components of the respiratory and central metabolic pathway although this is yet to be shown. We suspect that the potentially harmful mutations we report on will produce different growth phenotypes (Table 3). Therefore, we propose functional genomic assays followed subsequently by in vitro and in vivo characterization of mutants to confirm the contribution to growth and survival that each gene makes. Such studies on the MAF lineages are limited, but not for MTB, particularly the generalist MTB L4. Recent studies provide data describing functional consequences of synonymous SNPs (109–111), i.e. caution needs to be taken in inferring the relative significance or impact of observed genomic mutations from sequencing data alone. Therefore, further studies to correlate genotype with phenotype are necessary and our findings serve as a prelude to such experimental studies.

**Table 3:** Future proposed experiments.

| Examination | Proposed Experiments | Predicted Phenotype/<br>Expected outcome |
| --- | --- | --- |
| Experimentally determine if mutations detected in *eno*, *pca*, *pgi*, *pckA*, *korB*, *fba*, *aceE* and *glpK* of the Glycolytic Pathway and TCA cycle contribute to slow growth. | Mutagenesis experiments, complementation studies, Growth studies.<br>Mutagenesis studies would involve introducing the mutations detected into L2, L4 or H37Rv and testing for attenuated growth through growth studies or by complementing MAF and MTB L1 and L3 with a wild-type allele and testing for complementation of growth. | It is expected that the reported mutations will lead to attenuated growth. |
| Experimentally determine if mutations detected in *ndhA*, *pruB*, *qcrC*, *ctaB*, *frdAB*, *sdhA*, *atpDHG*, *ald*, *Rv0249c*, *narG* and *cydB* of the respiratory pathway contribute to slow growth. | Mutagenesis experiments, complementation studies, Growth studies.<br>Mutagenesis studies would involve introducing the mutations detected into L2, L4 or H37Rv and testing for attenuated growth in growth studies or by complementing MAF and MTB L1 and L3 with a wild-type allele and testing for | It is expected that the reported mutations will lead to attenuated growth. |
|  | complementation of growth. |  |
| Energy generation, in the form of ATP, for cellular processes | ATP quantitation assays. | MAF lineages will generate less ATP via the respiratory chain compared to MTB lineages |
| ROS accumulation and DNA damage in MAF cultures relative to MTB | Flow cytometry experiments and terminal deoxynucleotidyl transferase dUTP nick end labelling (TUNEL) assay. | Higher levels of ROS in MAF cultures and greater DNA damage of MAF relative to MTB. |
| Host-Pathogen interactions driving geographic restriction of MAF lineages to West Africa | Host genetics and Genome-wide association studies. | Specific genes in West Africans have undergone strong positive selection and increased the susceptibility of West Africans to MAF infection, permitting host restrictedness. |

#### 4.1.2 Investigate Bioenergetics in MAF lineages

It is highly plausible that adaptation of MAF lineages to growth under low oxygen (microaerobic) conditions could be a strategy to escape the harmful effects of ROS generated as electrons leak from a defective respiratory chain and react with built up oxygen (112). Moreover, reduced aerobic respiration or oxygen consumption in the MAF lineages could potentially affect the sensitivity of rapid diagnostics like the automated MGIT 960 system, that depends on oxygen consumption to detect growth (113). Interestingly, the percentage of MAF in certain parts of West and Central Africa was reported to have suddenly and sharply declined over the last decade (4, 114–117), which could possibly be due to the introduction and use of new diagnostics like the MGIT 960 system. Of note, we recently detected, in a retrospective analysis based on genotyping, that 84% of strains that did not grow in the MGIT 960 system were MAF L6 (Ofori-Anyinam *et al*., unpublished). Since growth and survival, driven by nutrient and energy metabolism, sustain pathogen transmission, and given that the success of drugs and phenotypic diagnostics rely on metabolic properties of the bacteria, understanding metabolism in these bacteria is essential. We hypothesize that MAF lineages generate less ATP and are more exposed to ROS than MTB lineages (Table 3). Therefore, we propose studies to investigate energy metabolism in MAF lineages relative to the MTB lineages, including whether MAF lineages are more exposed to ROS during growth.

#### 4.1.3 Host Genetics

One common feature of host specificity is genomic decay. Genomes of specialists usually show signs of genome decay evidenced by gene deletions or gene inactivation via point mutations. A driving force for the accumulation of mutations in some key electron transport and carbon metabolism genes in the specialist/host-restricted MAF lineages could indicate adaptation of the MAF lineages to a specific host, West Africans, or a niche within the host, due to altered or loss of function of these genes. This potential niche adaptation could be driven by a precise feature of the host environment that favors the association between MAF lineages and West Africans such as has been described for some diseases (118). Some findings have been made, including MAF lineage-specific mutations in genes and pseudogenes involved in vitamin B12 and vitamin B3 metabolism, important cofactor biosynthetic pathways for many cellular functions. Unlike MAF though, MTB is fully capable of synthesizing vitamin B12. Therefore, it has been suggested that the mutations in the vitamin B12 pathway of the MAF lineages may affect their host range to West Africans. According to a study in the United States, black persons reportedly have higher levels of vitamin B12 relative to other ethnicities (8, 10, 119–121). Future studies should investigate the molecular mechanisms underlying host specificity.

## 5 Supplementary Information

Supplementary File S1

Supplementary File S2

Supplementary File S3

Supplementary File S4

Supplementary File S5

## 6 Author Contributions

BO, FG, BdJ and CM designed the study. BO, FG, AR, TJ, BS and CM undertook analyses. BO, BdJ, FG and CM wrote the manuscript. All authors contributed discussion and reviewed the final manuscript.

## 9 Acknowledgments

The authors thank all study participants and colleagues for their contribution to the present analysis.

